# High-throughput genotype based population structure analysis of selected buffalo breeds

**DOI:** 10.1101/395681

**Authors:** Prakash B. Thakor, Ankit T. Hinsu, Dhruv R. Bhatiya, Tejas M. Shah, Nilesh Nayee, A Sudhakar, Chaitanya G. Joshi

## Abstract

The water buffalo (*Bubalus bubalis)* has shown enormous milk production potential in many Asian countries. India is considered as the home tract of some of the best buffalo breeds. However, genetic structure of the Indian river buffalo is poorly understood. Hence, for selection and breeding strategies, there is a need to characterize the populations and understand the genetic structure of various buffalo breeds. In this study, we have analysed genetic variability and population structure of seven buffalo breeds from their respective geographical regions using Axiom^®^ Buffalo Genotyping Array having 124,030 Single Nucleotide Polymorphisms (SNPs). Blood samples were obtained from 302 buffaloes comprising Murrah, Nili-Ravi, Mehsana, Jaffarabadi, Banni, Pandharpuri and Surti breeds. Diversity, as measured by expected heterozygosity (H_e_) ranged from 0.364 in the Surti to 0.384 in the Murrah breed. All the breeds showed negligible inbreeding coefficient. Pair-wise F_ST_ values revealed the lowest genetic distance between Mehsana and Nili-Ravi (0.0022) while highest between Surti and Pandharpuri (0.030). Principal component analysis and structure analysis unveiled the differentiation of Surti, Pandharpuri and Jaffarabadi in first two PCs, while remaining breeds were grouped together as a separate single cluster. Murrah and Mehsana showed early linkage disequilibrium decay while Surti breed showed late decay, similarly LD based Ne was drastically declined for Murrah and Mehsana since last 100 generations. In LD blocks to QTLs concordance analysis, 14.19 per cent of concordance was observed with 873 (out of 1144) LD blocks overlapped with 8912 (out of 67804) QTLs. Overall, total 4090 markers were identified from all LD blocks for six types of traits. Results of this study indicated that these SNP markers could differentiate phenotypically distinct breeds like Surti,Pandharpuri and Jaffarabadi but not others. So, there is a need to develop SNP chip based on SNP markers identified by sequence information of local breeds.

**Author Summary:** Indian buffaloes, through 13 recognised breeds, contribute about 49% in total milk production and play a vital role in enhancing the economic condition of Indian farmers. High density genotyping these breeds will allow us to study differences at the molecular level. Evolutionary relationship and phenotypes relations with genotype could be tested with high density genotyping. Breed structure analysis helps to take effective breeding policy decision. In the present study, we have used the high-throughput microarray based genotyping technology for SNP markers. These markers were used for breed differentiation using various genetic parameters. Population structure reflected the proportion of breed admixture among studied breeds. We have also tried to dig the markers associated with traits based LD calculation. However, these SNPs couldn’t explain obvious variation up to the expected level, hence, there is need to develop an indigenous SNP chip based on Indian buffalo populations.

## Introduction

The importance of genetic diversity in livestock is directly related to the need for genetic improvement of economically important traits as well as to facilitate rapid adaptation to potential changes as per breeding goals [1]. Population structure, and unusual levels of shared ancestry, can potentially cause spurious associations. The analysis of a large number of SNPs across the genome will reveal aspects of the population genetic structure, including evidence of adaptive selection across the genome [2]. Domestication greatly changed the morphological, behavioural characteristics, and selection programmes for improving the production traits allowed the formation of very diverse breeds [3].

India, the largest producer of milk in the world, is producing over 155.5 million tone milk during 2015-16 and about 49% of milk production is contributed by buffaloes [4]. India has approximately 108.7 million buffaloes [4] with 13 registered breeds recognized based on their phenotypic characteristics, production performance, utility pattern and eco-geographical distribution.

Genetic analysis is facilitated by genotyping polymorphic genetic loci, also called genetic variants, signspots, landmarks or markers. SNPs are the most common type of genetic variants, consisting of a single nucleotide differences between two individuals at a particular site in the DNA sequence. SNPs are generally bi-allelic. Assessing genetic biodiversity and population structure of minor breeds through the information provided by neutral molecular markers like, SNPs & microsatellites, allows determination of their extinction risk and to design strategies for their management and conservation [5]. Maintenance of genetic variation is a condition for continuous genetic improvement. For overall breed improvement and to meet future challenges there is an immediate action to be taken for characterization of buffalo breeds in India. Comprehensive knowledge of genetic variation within and among different breeds is very much necessary for understanding and improving traits of economic importance. Current study was performed based on SNP genotyping data to determine the genetic structure of Indian buffalo breeds so that to construct appropriate conservation strategies and to utilize the breed variation.

## Materials and Methods

### Animals and Sampling

A total of 302 female buffaloes were used in this study, comprising of seven breeds: Murrah (*n*=70), Nili-Ravi (*n* = 40), Mehsana (*n* = 75), Jaffarabadi (*n* = 41), Banni (*n* = 20), Pandharpuri (*n* = 34) and Surti (*n* = 22). All animals were selected based on their true breed specific phenotypic characteristics from their respective home tract and blood samples were collected from all the selected animals.

### SNP Genotyping

DNA was extracted using QIAamp^®^ kit as per manufacturer’s instructions at R&D laboratory NDDB, Hyderabad. DNA quantity and quality were checked using Nanodrop^™^ (Thermo Fisher Scientific, MA) and agarose gel electrophoresis respectively. SNP genotyping was carried out using Axiom^®^ Buffalo Genotyping Array with 123,040 SNPs on GeneTitan^®^ MC (Thermo Fisher Scientific, MA) instrument at a commercial laboratory (Imperial Life Science Group, Gurgaon). Array was pre-designed through the Expert Design Program, facilitated by Affymetrix and developed in collaboration with the International Buffalo Genome Consortium using reference genome of *Bos taurus* (UMD3.1) for SNP position and annotation (Thermo Fisher Scientific, MA; Iamartino *et al.*, 2013). It was designed based on SNPs discovered from Mediterranean, Murrah, Jaffarabadi and Nili-Ravi breeds of buffaloes. The genotyping experiment was performed in four batches, NDDB_EXP 1 (96 samples), NDDB_EXP 2 (96 samples), NDDB_EXP 3 (95 samples) and NDDB_EXP 4 (89 samples) with average call rate ranged from 97 per cent to 98.8 per cent.

### Data filtering and quality control

Only SNPs mapped to autosomal chromosomes were used in this study. Data was filtered based on criteria: SNPs that have poor call rate (<95%). Further, quality control was performed with PLINK v1.07 [6] and SNPs removed with following criteria: missing genotypes (geno < 0.1), individual missing genotypes (mind < 0.1), minor allele frequency (MAF < 0.05) and Hardy-Weinberg Equilibrium (HWE < 0.00001). Remaining markers were used for further analysis.

### Genetic Diversity Assessment

Observed and expected genotype frequencies within each breed was calculated for all the loci using PLINK v1.07 [7] and the results were evaluated based on p values obtained for each loci. Linkage disequilibrium was calculated using PLINK and R^2^ values were calculated for all SNP pairs which were located not more than 1000 SNPs apart and falling under 10 Mb distance windows. Further SNPs were binned with bin size of 10,000 bases distance and average R^2^ value of each bin was plotted against median distance value ggplot2 v2.2.1 [8] package in R v3.3. Pair-wise F^ST^ values between all possible combination of breeds were estimated and subsequently dendrogram was generated in Fitch-Phylip [9] using Fitch-Margoliash method.

Breed-wise effective population size (Ne) was calculated using SNeP v1.1 [10] with parameters: bin-width=50,000 bp; minimum distance between SNPs=50,000 bp, maximum distance between SNPs=4,000,000 bp, minimum allele frequency=0.05. Principle component analysis was calculated using PLINK-1.9 [11] with 285 highly variable markers (Allele frequency difference between breeds > 0.5). PCA was plotted using scatterplot3d [12] package in R. Breed structure and breed differentiation was performed using fastSTRUCTURE [13] using same 285 highly variable markers. The differentiation of populations was performed up to the group (K) level of 8 using simple model. The fastSTRUCTURE analysis provided ancestry proportions for each sample under analysis which was graphically represented by distruct.py script within the fastSTRUCTURE software.

### Genome wide LD block mapping on QTLs

Linkage disequilibrium (LD) blocks, combination of alleles linked along a chromosome and inherited together from a common ancestor, were generated with Java based gPLINK v1.0 and Haploview v2.01 [14]. Blocks were defined by employing haplotypic diversity criterion, where a small number of common haplotypes provide high chromosomal frequency coverage [15-18]. The algorithm suggested by Gabriel et al. [19] was used which defines a pair of SNPs to be in strong LD if the upper 95% confidence bound of D′ value between 0.7 and 0.98. Reconstructed haplotypes were inserted into Haploview v2.01 [14] to estimate LD statistics and construct the blocking pattern for all 29 autosomes. LD blocks were estimated using an accelerated EM algorithm method described by Qin et al. [20]. QTL database was retrieved from previously reported QTLs in Animal QTLdb [21]. QTL data set of cattle (*Bos taurus*) QTL_UMD_3.11.bed was used as a reference for the analysis, containing the information regarding six types of the traits: milk traits; health traits; production traits; reproduction traits; exterior traits; and meat and carcass traits. The QTL files were intersected with the files of LD-blocks using Bedtools v2.26.0 [22] to obtain information of QTLs overlapping with LD blocks.

## Results

### Genetic Diversity Analysis

After data filtering and quality filtering, 295 samples with 75,704 SNPs remained available for population analysis. SNPs were discarded (total 47,336 SNPs) based on criteria: poor quality call rate (42,166), unknown chromosome-specific position (17), Chromosome X (4228), HWE less than 0.00001 (528), missing genotype rate less than 0.1 (471), and all genotypes from seven Nili-Ravi animals were removed since they were outliers.

Alternate allele frequency followed almost intermediate distribution with higher proportion for Murrah and Mehsana (Fig 1.A). Highest allele count was observed in the range of frequency class 0.2-0.5. Highest average alternate allele frequency was observed in Nili-Ravi (0.3051) followed by Murrah (0.3049) while Jaffarabadi showed least average (0.3028) among all breeds (Fig 1.B). Highest proportion of alternate alleles was observed in Murrah with 91.86 per cent while lowest proportion was observed in Surti with 89.86 per cent (Fig 1.C). The observed heterozygosity (Ho) and expected heterozygosity (He) was also found highest in Murrah breed (0.3864 and 0.3846) followed by Mehsana breed (0.3857 and 0.3830), while lowest was observed in Pandharpuri breed with Ho = 0.3719 and He = 0.3680 (**Error! Reference source not found.** 1). The lowest F_IS_ were observed for Murrah (−0.0046) and Mehsana (−0.0070) while highest was seen in Surti (−0.0314) followed by Banni (−0.0270).

**Fig 1:**
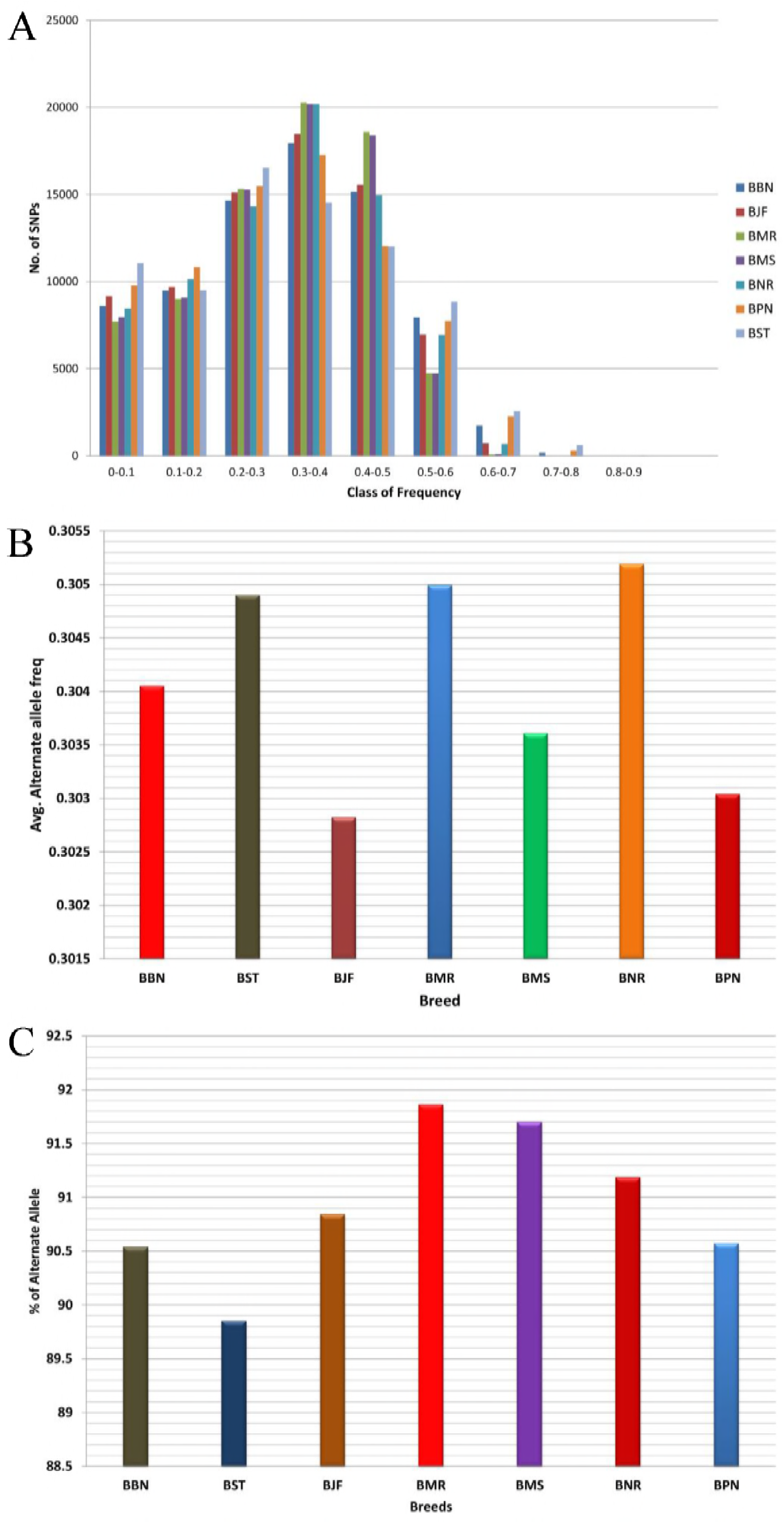
Alternate allele distribution. (A) Distribution of alternate allele frequency in studied buffalo breed (B) Breed-wise average alternate allele frequency distribution (C) Breed wise proportion and distribution of alternate allele with allele frequency > 0 (SNPs removed which are monomorphic)(BBN: Banni, BJF: Jaffarabadi, BMR: Murrah, BNR: Nili-Ravi, BMS: Mehsana, BPN: Pandharpuri, BST: Surti)

F_ST_ values showed lowest genetic distance between Murrah and Nili-Ravi (0.00221) followed by Murrah and Mehsana (0.00402) while highest genetic distance was observed between Surti and Pandharpuri (0.03097) followed by Surti and Banni (0.02650) (Table 2). Based on F_ST_ values, phylogenetic tree placed Nili-Ravi and Murrah as well as Mehsana and Banni together in two separate clusters, which corresponds with their geographical origin (Fig 2). This differentiation also correlates with the phenotypic differentiation of the buffalo breeds.

**Table 1:**
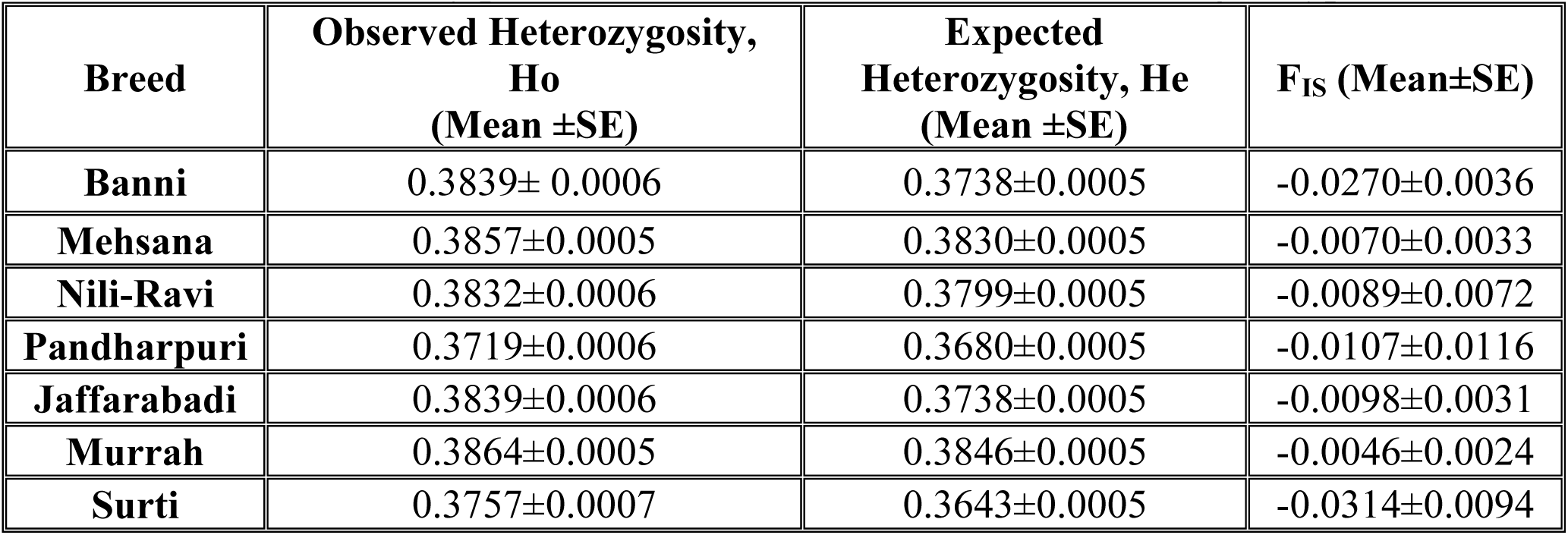
Genetic diversity parameters in Indian buffalo breeds from genotyped data

**Table 2:** Standard genetic distance or Mean pairwise F_ST_ values among various buffalo breeds

| **Breed** | **Murrah** | **Nili-Ravi** | **Mehsana** | **Jaffarabadi** | **Banni** | **Pandharpuri** | **Surti** |
| --- | --- | --- | --- | --- | --- | --- | --- |
| **Murrah** | 0 |  |  |  |  |  |  |
| **Nili-Ravi** | 0.00221 | 0 |  |  |  |  |  |
| **Mehsana** | 0.00402 | 0.00599 | 0 |  |  |  |  |
| **Jaffarabadi** | 0.00947 | 0.01209 | 0.01794 | 0 |  |  |  |
| **Banni** | 0.02143 | 0.00790 | 0.00442 | 0.01322 | 0 |  |  |
| **Pandharpuri** | 0.01833 | 0.02330 | 0.02188 | 0.02156 | 0.02650 | 0 |  |
| **Surti** | 0.02143 | 0.02430 | 0.01794 | 0.02122 | 0.02650 | 0.03097 | 0 |

**Table 3:** Chromosome wise LD block distribution statistics with total number of LD blocks, average block size, mean and maximum number of SNPs in blocks

| Chromosome | Total LD blocks | Mean number of SNPs per block | Max. Number of SNPs in blocks |
| --- | --- | --- | --- |
| 1 | 99 | 3.48 | 7 |
| 2 | 87 | 3.68 | 9 |
| 3 | 59 | 3.25 | 6 |
| 4 | 58 | 3.44 | 8 |
| 5 | 63 | 3.73 | 15 |
| 6 | 43 | 3.72 | 9 |
| 7 | 44 | 3.72 | 15 |
| 8 | 52 | 3.75 | 10 |
| 9 | 39 | 4.00 | 8 |
| 10 | 36 | 3.94 | 6 |
| 11 | 54 | 3.51 | 9 |
| 12 | 37 | 3.75 | 9 |
| 13 | 38 | 3.34 | 9 |
| 14 | 31 | 2.93 | 13 |
| 15 | 33 | 3 | 6 |
| 16 | 44 | 3.56 | 12 |
| 17 | 30 | 3.83 | 16 |
| 18 | 24 | 3.04 | 5 |
| 19 | 31 | 4.54 | 11 |
| 20 | 23 | 3.47 | 9 |
| 21 | 36 | 3.94 | 11 |
| 22 | 26 | 3.76 | 13 |
| 23 | 22 | 3.72 | 7 |
| 24 | 27 | 2.96 | 7 |
| 25 | 29 | 2.75 | 9 |
| 26 | 16 | 3.56 | 8 |
| 27 | 23 | 3.34 | 8 |
| 28 | 19 | 3.84 | 7 |
| 29 | 22 | 3.77 | 10 |
| All | 1145 | 3.56 |  |

**Fig 2:**
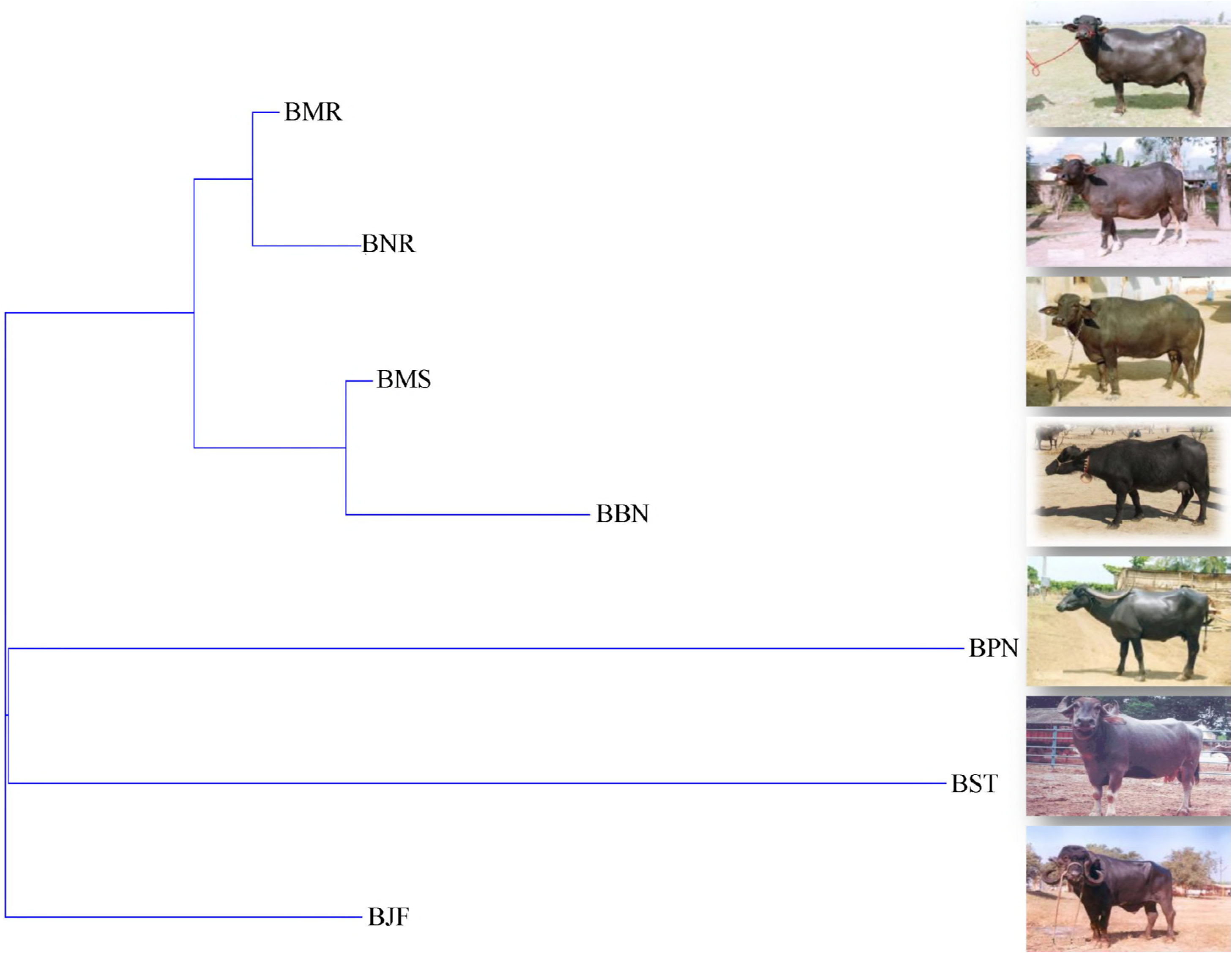
Dendrogram of breed differentiation based on pair-wise F_ST_ values. Labelled tree with name of breed at each leaf (BBN: Banni, BJF: Jaffarabadi, BMR: Murrah, BNR: Nili-Ravi, BMS: Mehsana, BPN: Pandharpuri, BST: Surti)

### Population Structure

The total variability of principal components explained was 65.6 per cent of which by first, second and third components explained 30.05 per cent, 27.14 per cent and 8.45 per cent, respectively. This variation resulted in separate cluster of Surti, Pandharpuri and Jaffarabadi on coordinates 1, 2 and 3 respectively while other breeds remain admixed (Fig 3).

**Fig 3:**
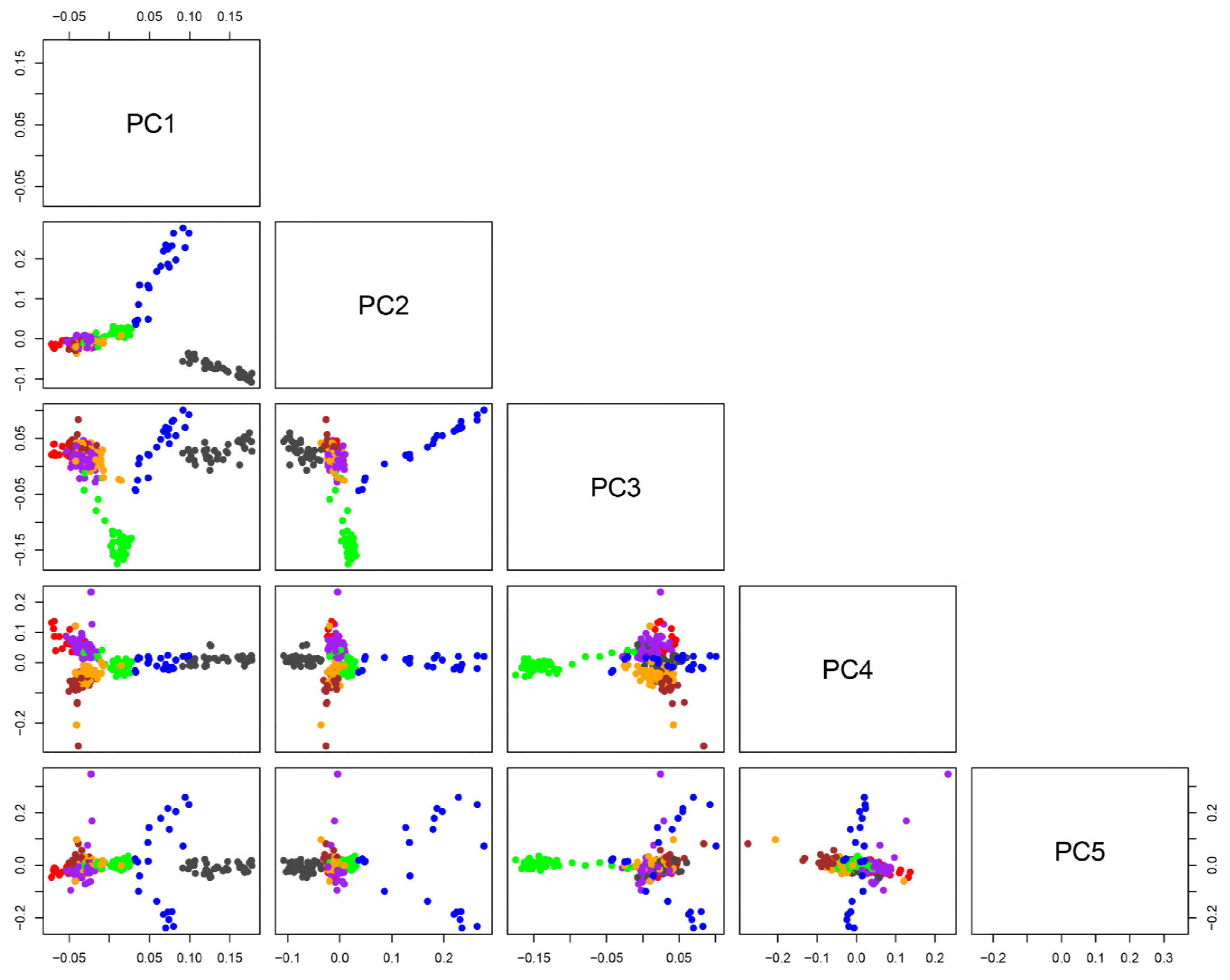
2D PCA plot of all seven buffalo breeds together up to principal components 5. (BBN: Banni, BJF: Jaffarabadi, BMR: Murrah, BNR: Nili-Ravi, BMS: Mehsana, BPN: Pandharpuri, BST: Surti)

Further, relatedness between breeds and the significance of the existence of subpopulations was investigated by model-based unsurprised clustering using K=2 to K=8 (K values indicates the number of groups). Banni breed showed better separation with small amount of admixture at all levels while Murrah and Mehsana breed showed higher amount of admixture consistent with its crossing with other breeds. With increasing K values, Pandharpuri and Surti showed separation at all subsequent levels (Fig 4). At K=7, four buffalo breeds (Surti, Pandharpuri, Jaffarabadi and Banni) were distinctly separated. Three Jaffarabadi breed were identified as pure breed based on Q-value greater than 95 per cent while remaining showed variable amount of admixture. Similarly, Pandharpuri buffaloes showed highest number (26) of purebred individuals. Likewise, Surti breed showed negligible admixture with other breeds.

**Fig 4:**
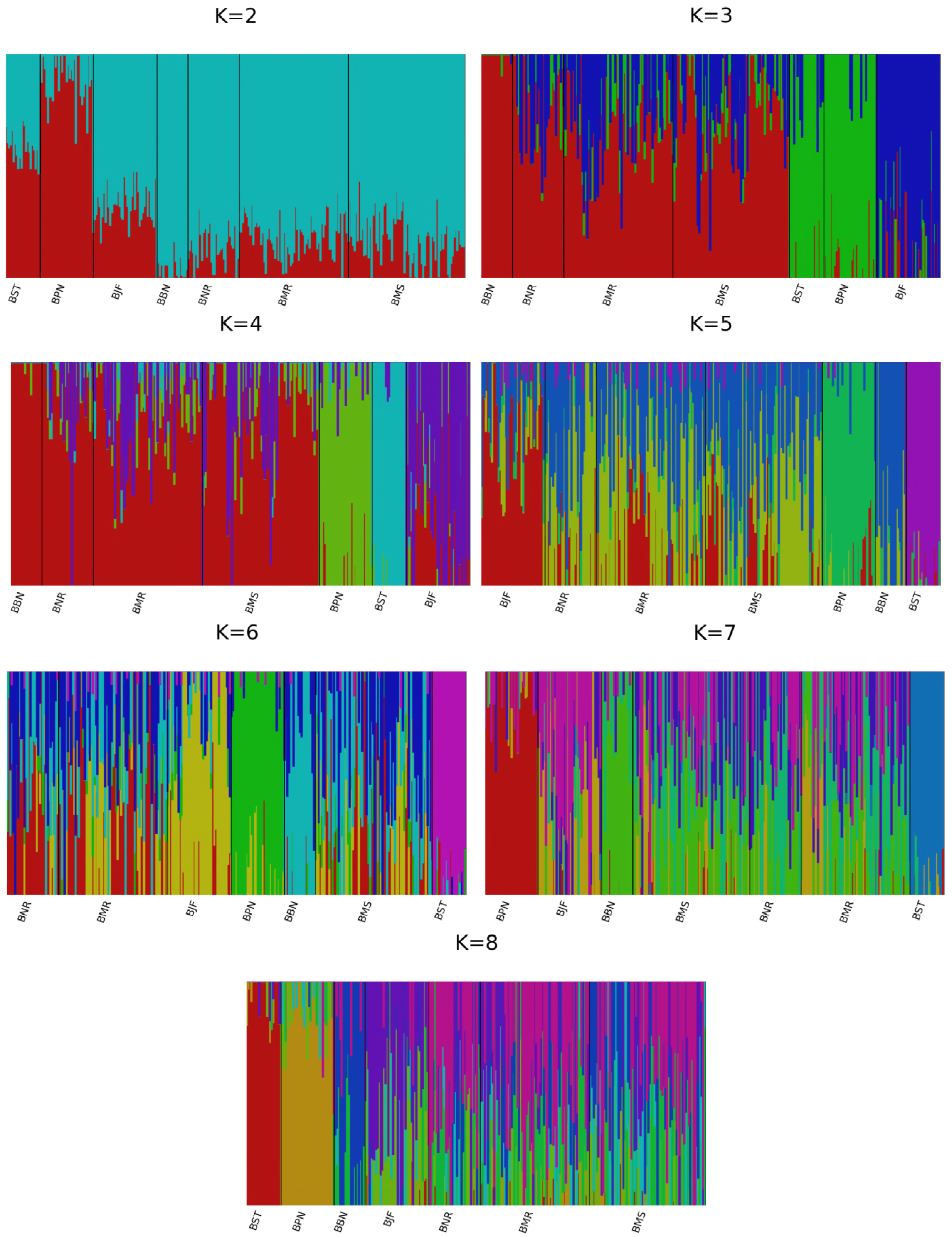
Estimated population structure by fastSTRUCTURE for K = 2 to K = 8. Each individual is represented by a thin vertical line, and each breed is demarcated by a thick vertical black line. (BBN: Banni, BJF: Jaffarabadi, BMR: Murrah, BNR: Nili-Ravi, BMS: Mehsana, BPN: Pandharpuri, BST: Surti)

### Linkage Disequilibrium Analysis

LD decay was performed using bin size of 10 kb distance between SNPs. LD decay showed highest R^2^ value in Surti (from 0.412 to 0.175) followed by Banni (from 0.412 to 0.169). While Pandharpuri (from 0.379 to 0.149) and Nili-Ravi (from 0.412 to 0.139) as well as Mehsana (from 0.378 to 0.128) and Murrah (0.382 to 0.120) decayed almost with same rate. In Surti breed, LD decayed late as distance between loci increased compared to other. Nili-Ravi and Pandharpuri decayed almost together with given distance. Similar trend was shown by Mehsana and Murrah. Moreover, Mehsana and Murrah showed early decay among all the breeds (Fig 5. A).

**Fig 5:**
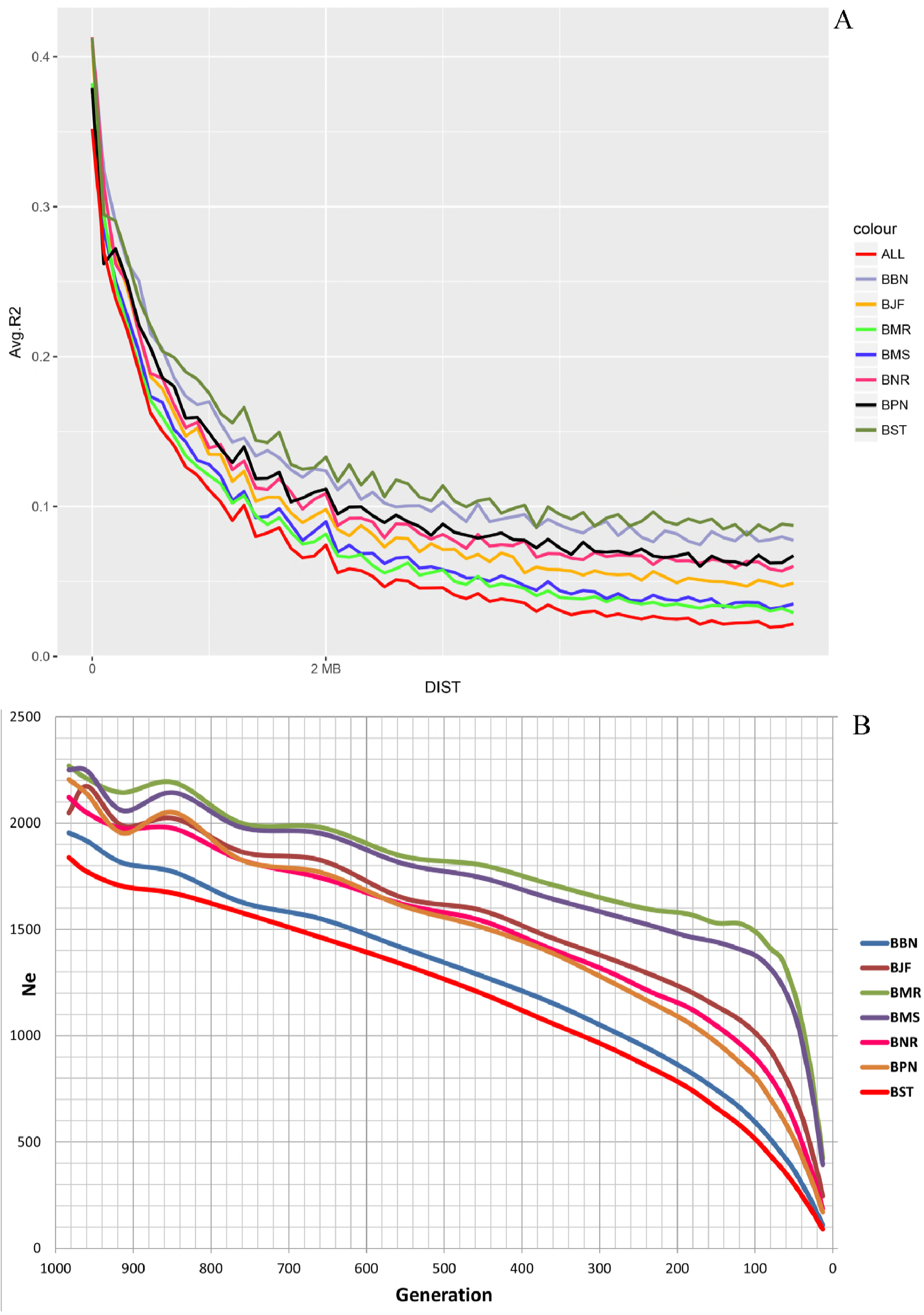
Linkage Disequilibrium study of Buffalo breeds: (A) Linkage disequilibrium (LD) decay plot based on all pairwise comparisons between adjacent loci of all seven breeds. The horizontal axis depicts the intermarker distance in base pair and vertical axis shows the average R^2^ values (B) **Effective population size (Ne) of different breeds with respect to generation time** (BBN: Banni, BJF: Jaffarabadi, BMR: Murrah, BNR: Nili-Ravi, BMS: Mehsana, BPN: Pandharpuri, BST: Surti)

A continuous steady decline in effective population size was observed over last 1000 generations in all breeds. Effective population size of Murrah and Mehsana has drastically declined over last 100 generations with an increasingly steeper slope while Surti and Banni are declining almost at constant rate (Fig 5.B). Jaffarabadi, Nili-Ravi and Pandharpuri showed intermediate rate of declination over last 100 generations.

### Genome-Wide Study of LD blocks

#### LD blocks

Total 1144 LD blocks were obtained with highest number of blocks on chromosome 1 (99 blocks) while lowest number of blocks on chromosome 28 (19 blocks) (Error! Reference source not found.). Overall, mean number of SNPs in block ranged from 2.75 to 4.54 SNPs per chromosome while, maximum number of SNPs per block ranged from 5 (chromosome 18) to 16 (chromosome 17). Overall, frequency-based size distribution of LD blocks revealed that highest number (547) of LD blocks were found having size less than 50 kb while very few (8) were observed having size as high as 400-450 kb (Fig 6).

**Fig 6:**
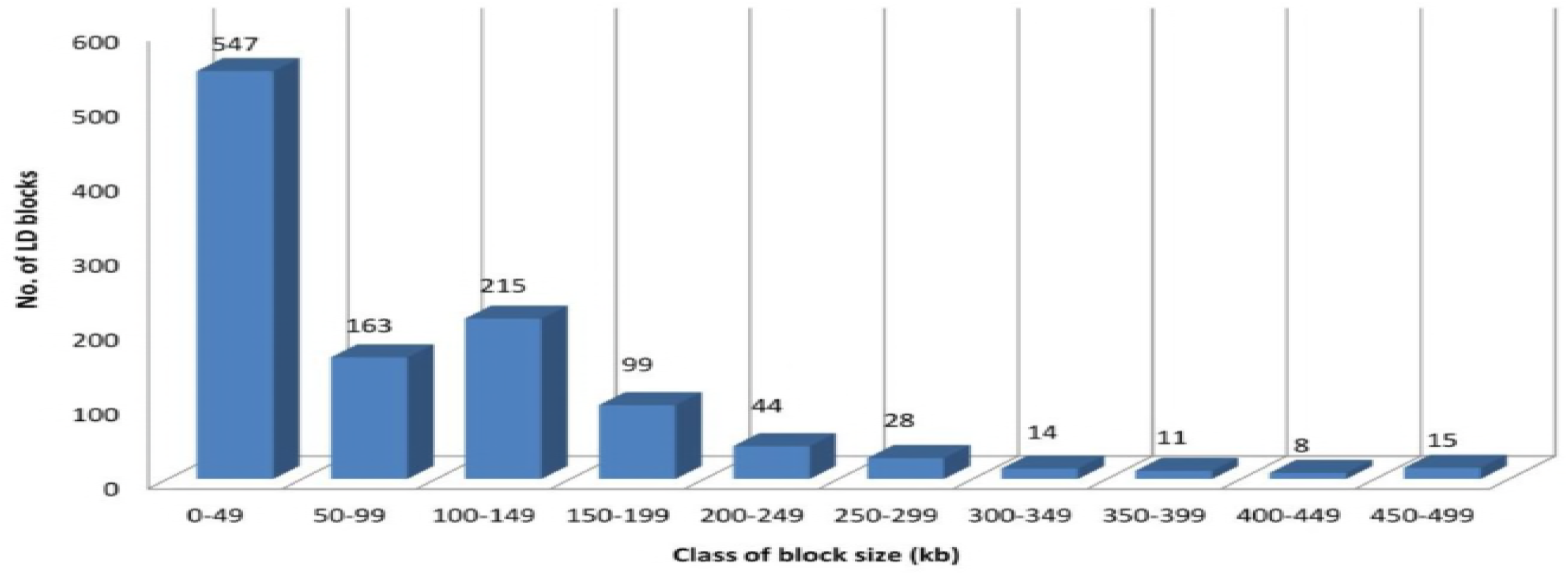
LD blocks distribution based on size of block in respective class of size (in kb)

#### LD blocks – QTL concordance

Out of 1144 LD block (4090 markers), 436 LD blocks (1624 markers), 368 LD blocks (1285 markers), 326 LD blocks (1253 markers),345 LD blocks (1351 markers), 81 LD blocks (338 markers) and 104 LD blocks (426 markers) overlapped with QTLs for milk production trait; meat and carcass trait; reproduction trait; production trait; exterior trait; and health trait respectively (Fig 7). Concordance, measured as proportion of LD blocks and QTLs overlapping each other, was highest in chromosome one (16.91 %) while lowest on chromosome 14 (0.91 %). Overall concordance of all the chromosomes together was 14.19%, with 873 LD blocks intersecting with 8947 QTLs (Table 4). Chromosome-wise distribution of LD-blocks, number of markers and mapped QTLs for respective traits is shown in S1 Table.

**Table 4:**
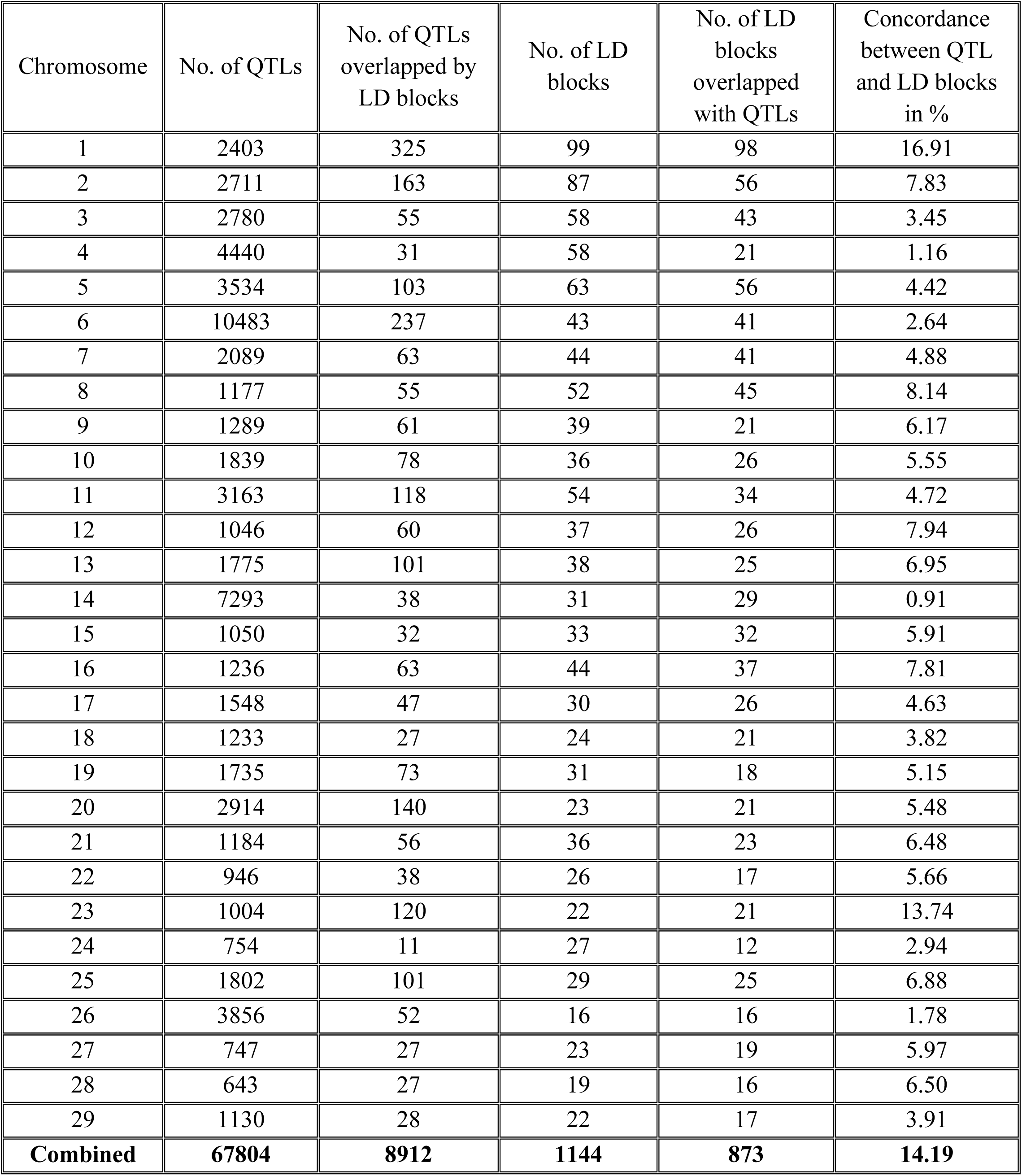
Chromosome-wise distribution of LD blocks and QTLs with its percentage of concordance and discordance

| Chromosome | No. of QTLs | No. of QTLs overlapped by LD blocks | No. of LD blocks | No. of LD blocks overlapped with QTLs | Concordance between QTL and LD blocks in % |
| --- | --- | --- | --- | --- | --- |
| 1 | 2403 | 325 | 99 | 98 | 16.91 |
| 2 | 2711 | 163 | 87 | 56 | 7.83 |
| 3 | 2780 | 55 | 58 | 43 | 3.45 |
| 4 | 4440 | 31 | 58 | 21 | 1.16 |
| 5 | 3534 | 103 | 63 | 56 | 4.42 |
| 6 | 10483 | 237 | 43 | 41 | 2.64 |
| 7 | 2089 | 63 | 44 | 41 | 4.88 |
| 8 | 1177 | 55 | 52 | 45 | 8.14 |
| 9 | 1289 | 61 | 39 | 21 | 6.17 |
| 10 | 1839 | 78 | 36 | 26 | 5.55 |
| 11 | 3163 | 118 | 54 | 34 | 4.72 |
| 12 | 1046 | 60 | 37 | 26 | 7.94 |
| 13 | 1775 | 101 | 38 | 25 | 6.95 |
| 14 | 7293 | 38 | 31 | 29 | 0.91 |
| 15 | 1050 | 32 | 33 | 32 | 5.91 |
| 16 | 1236 | 63 | 44 | 37 | 7.81 |
| 17 | 1548 | 47 | 30 | 26 | 4.63 |
| 18 | 1233 | 27 | 24 | 21 | 3.82 |
| 19 | 1735 | 73 | 31 | 18 | 5.15 |
| 20 | 2914 | 140 | 23 | 21 | 5.48 |
| 21 | 1184 | 56 | 36 | 23 | 6.48 |
| 22 | 946 | 38 | 26 | 17 | 5.66 |
| 23 | 1004 | 120 | 22 | 21 | 13.74 |
| 24 | 754 | 11 | 27 | 12 | 2.94 |
| 25 | 1802 | 101 | 29 | 25 | 6.88 |
| 26 | 3856 | 52 | 16 | 16 | 1.78 |
| 27 | 747 | 27 | 23 | 19 | 5.97 |
| 28 | 643 | 27 | 19 | 16 | 6.50 |
| 29 | 1130 | 28 | 22 | 17 | 3.91 |
| **Combined** | **67804** | **8912** | **1144** | **873** | **14.19** |

**Fig 7:**
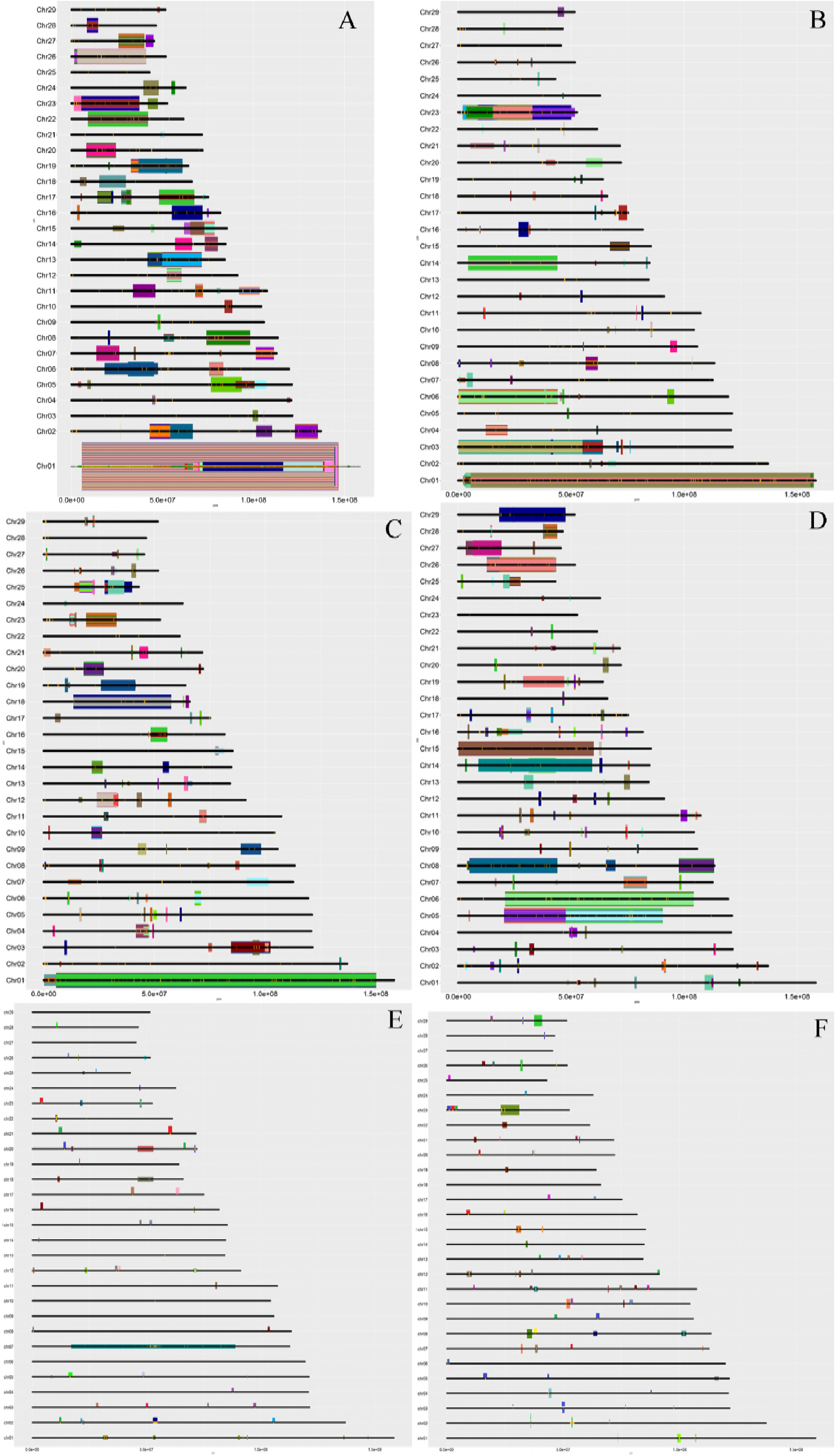
Concordance of LD blocks with QTLs (A) Milk production traits (B) Production traits (C) Reproduction traits (D) Meat and carcass traits (E) Health trait and (F) Exterior traits. Vertical axis shows the chromosome number, horizontal axis shows the base pair position, thick middle black bar shows physical length of chromosome, thin orange colored bars over black bars shows LD blocks and the colored segments reflects the physical length of QTLs.

Further, dendrogram was plotted based on markers overlapping with milk fat percentage (143 markers) and body weight (315 markers) QTLs (Fig 8). Surprisingly, no pattern was observed linking phenotypic recorded data with marker-based separation.

**Fig 8:**
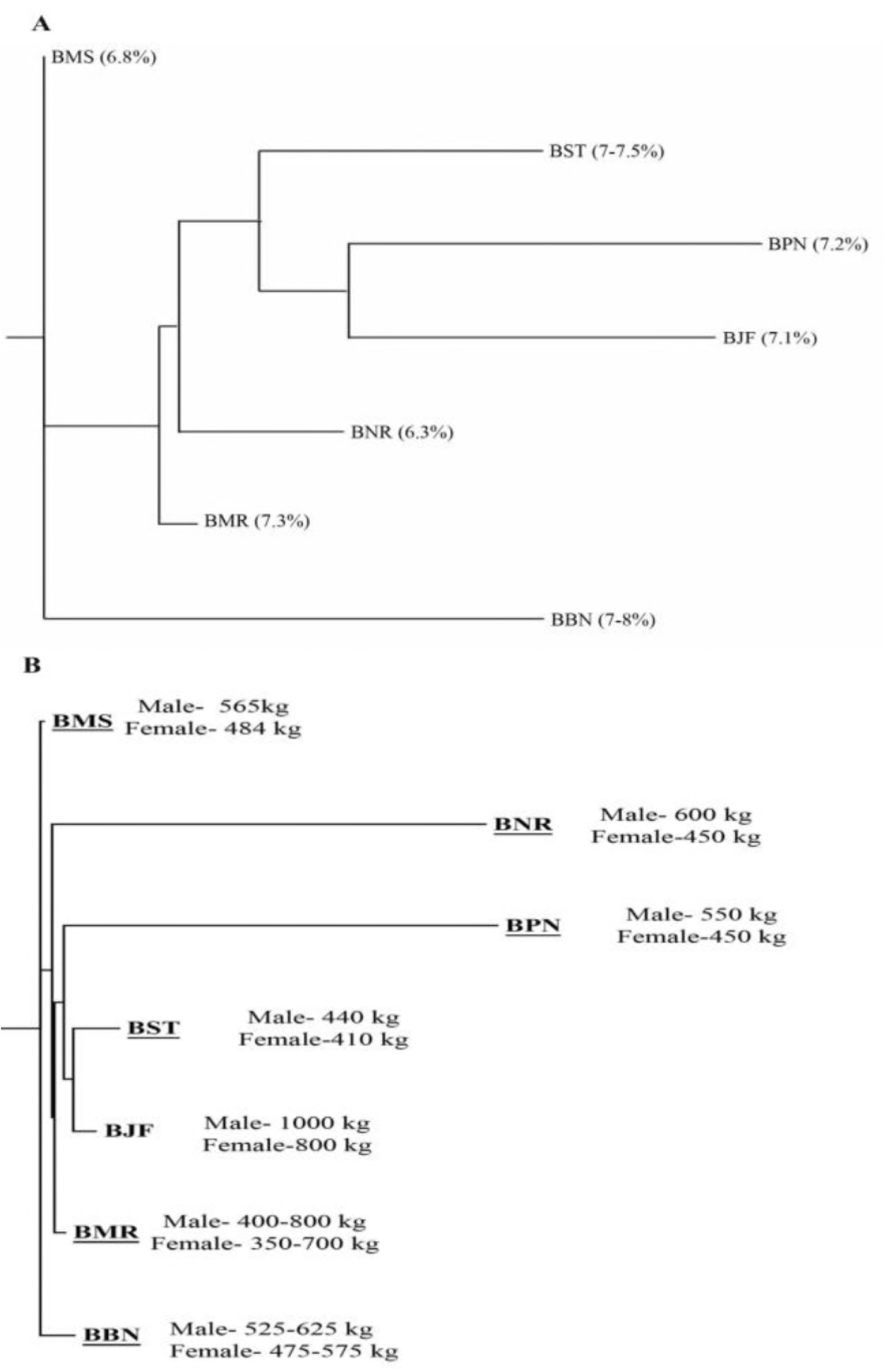
Trait based dendrogram of studied buffalo breeds (A) Dendrogram of studied buffalo breeds based on markers covered by fat percentage QTLs (Fat percentage was sourced from INAPH data, NDDB and ICAR) (B) Dendrogram of studied buffalo breeds based on markers covered by body weight QTLs (Body weight was sourced from ICAR) (BBN: Banni, BJF: Jaffarabadi, BMR: Murrah, BNR: Nili-Ravi, BMS: Mehsana, BPN: Pandharpuri, BST: Surti)

## Discussion

Genetic diversity studies conducted for buffalo in India have previously relied primarily on the use of microsatellites markers [23-28] while use of SNP genotype data in Indian cattle has been previously reported by Dash et al. [29].

The chip used in this study was designed based on SNP markers of 4 breeds (Mediterranean, Murrah, Nili-Ravi and Jaffarabadi) although using the reference of *Bos taurus* (UMD_3.1 assembly) [30]. The differences in allele frequencies among the breeds may be caused by genetic drift, adaptation to selection or ancient divergence among founder populations [31,32]. Therefore, it is possible that the SNPs that have been identified as being useful in one population may not necessarily be as useful in other breeds. Here, we used the term ‘Alternate allele’, because minor allele frequency does not exceed over 0.5 while in this study, the allele frequency exceeds over 0.5 often called as ‘Fixed allele’ and hence, it has been considered as an “Alternate allele’. The differences in observed allele frequencies among breeds show the genetic diversity that exists within and between the breeds [33]. The overall allele frequency observed in this study was higher than previously reported studies in indicine breeds [34-36].

Murrah and Mehsana had the highest numbers SNPs with intermediate class of frequency suggesting that this array could be utilised for these breeds for association studies, with available phenotypic data for the traits of interest. The higher genetic variability observed in the Murrah and Mehsana, which is evident from the population structure analysis that suggests introgression of these breeds with other breeds such as Banni, Nili-Ravi, Jaffarabadi, etc. While Surti and Pandharpuri showed less polymorphic SNPs suggesting less genetic variability. These findings further supported by observed heterozygosity (Ho) and expected heterozygosity (He) values, which was found higher in Murrah and Mehsana breeds as compared to other breeds which could be due to extensive use of these two breeds via artificial insemination technique. The purpose of using these breeds is to obtain appropriate production since they are the good milk producers. Pandharpuri and Surti have less genetic variability with the lowest He suggesting that inbreeding in conjunction with a small population size and resulted in a loss of variation within the breed. This low diversity was previously reported in other studies of cattle and buffalo using microsatellites [37-39] and using SNP panels [29,40]. The F statistics is an estimate of variation due to differences among populations, which is the reduction in heterozygosity of a sub-population due to genetic drift. All breeds have shown negligible inbreeding as negative values of F_IS_ in all breeds indicate that there is absence of inbreeding in these breeds. In this study, the mean F_ST_ indicated that a pair of Surti and Pandharpuri population has greater genetic distance than other pairs, similar to results of European cattle breeds (Brown Swiss and Holstein Friesian) [40]. Phylogenetic tree based on F_ST_ values revealed that grouping was observed according to geographical distribution of population as shown in microsatellite based study of cattle performed by Shah et al. [41]. They displayed results of phylogenetic relationships as three main clusters according to geographical distribution: Dangi and Khillar (cluster I); Gir, Kankrej, Nimari and Malvi (cluster II); and Gaolao and Kenkatha (cluster III). However, the results failed to explain the hypothesis that Mehsana breed has been developed using Murrah bulls on local Surti buffaloes [28] as both the breeds were clustered separately. In case of genetic diversity (F_ST_) of buffalo based on microsatellite markers [42], similar cluster pattern was observed as in current study. Surti and Pandharpuri grouped in single cluster in present study as shown by Kumar *et al.* (2007) as; cluster of Mehsana with Jaffarabadi, Surti with Pandharpuri and Murrah with Nagpuri. However, Jaffarabadi and Mehsana grouped in different clusters in present study whereas they were grouped in single cluster in the study updated Kumar *et. al.* (2007).

The results of the PCA analysis revealed the higher amount of genetic similarities among Murrah, Mehsana, Banni and Nili-Ravi, while Surti, Jaffarabadi and Pandharpuri showed greater genetic differentiations with three distinct clusters. The clustering of populations from both the PCA and fastSTRUCTURE indicated low levels of within population diversity of the Surti, Jaffarabadi and Pandharpuri breeds and higher divergences of these populations from the Murrah, Mehsana, Banni and Nili-Ravi breeds. In current study, Surti, Jaffarabadi and Pandharpuri grouped in separate clusters, however, it was shown in single cluster by Kumar et al. [25]. The high genetic diversity and distinct breed structure imply the possibility of selective breeding in these Indian buffalo breeds for genetic improvement (Murrah and Mehsana). Four breeds (Surti, Pandharpuri, Jaffarabadi and Banni) were able to get distinctly separate while two breeds (Murrah and Mehsana) showed greater admixture. These two breeds have been most popular amongst the buffalo breeds in terms of high milk yield. Murrah semen has been extensively and indiscriminately used for artificial insemination (AI) across the country while Banni, Jaffarabadi and Pandharpuri are less in number and been less utilized for insemination throughout the country, which has led to a steady decline in the genetic diversity present in the non-descript or less characterized populations. Kumar et al. [25] evaluated the breed admixture using microsatellite markers and results revealed that the 3 different clusters contributed mainly from the Toda, Jaffarabadi and Pandharpuri animals, with a very high membership coefficient. In case of cattle using microsatellite markers [41], the differentiation of Dangi, Khillar and Kenkatha cattle breeds was performed while Kankrej showed greater admixture with other breeds.

The probable cause of drastic decline is too large distribution of population from which only small proportion of population of superior germplasm being used for breeding purpose through AI. Moreover, in past, before 100-150 generations, farmers had adapted the intensive selective breeding based on some characters and use of elite animals from certain areas in absence of AI. Murrah has higher average allele frequencies while Pandharpuri and Surti breeds has lower values can be interpreted as higher allele frequency can be ascertained biasness to SNP selection from Murrah reference.

LD decay used to study the linkage of markers with increase in intermarker distance and was to decide appropriate intermarker distance for different populations. The magnitude of LD and its decay with genetic distance determine the resolution of association mapping and are useful for assessing the desired numbers of SNPs on arrays. The results of LD decay illustrate Surti breed showing early decay as compared to other breeds while Mehsana and Murrah breeds showed late decay together which could be assumed as they are under strong selection pressure. Similar results were obtained by Dash et al. [29] for Indian cattle breeds where Sahiwal and Tharparkar breeds showed late decay. These results reflected that the Surti breed has smaller population size as it got decayed earlier. Other breeds also exhibited LD decay as per their available breedable population. Larger the population size, longer the LD decay. Effective population size of Murrah and Mehsana has drastically declined over last 100 generations. It is believed that Mehsana breed has been developed a couple of centuries ago from Murrah and Surti buffalo (might have completed less than 100 generations). Hence, the results should be viewed in light of theoretical expectations. It gives information regarding effective population size of ancestors. Shin et al. [43] estimated the effective population size in Korean cattle which revealed rapid increase in effective population size over the past 10 generations with the values increasing fivefold (close to 500) by 10 generations. Santana et al. [44] also reported small effective size (40) from several Murrah herds. An effective population size of at least 50 animals is enough to prevent inbreeding depression, the minimum level recommended by the FAO (2007).

The haplotype block structure and its distribution in the genome of cattle, especially studies based on high density SNPs, have been rarely reported [45]. Thus, the current analysis was performed to construct the haplotype structure in the buffalo genome and to detect the relevant genes affecting quantitative traits. Jiang et al. [46] identified the milk trait QTL specific SNPs in cattle and found a large proportion of the significant SNPs (61 out of 105) were located on BTA14 and that were also located within the reported QTL regions. In our study, 76 QTLs (mostly of milk protein percentage, milk yield and milk fat per cents) on chromosome 20 concordant with 13 LD blocks. Mai et al. [47] recognized total 98 QTLs for milk production trait, which included 30 for milk index, 50 for fat index, and 18 for protein index. The density of QTLs of body weight was higher on chromosome 23 along with other productive traits. Mai et al. [47] reported a greater number of significant SNPs associations for production (54) than for fertility traits (29) with 22 QTL regions associated with fertility traits and 14 with production traits. The concordance study of meat and carcass trait revealed that the largest QTL of shear force was observed on chromosome 6 and QTL of tridecylic acid content located on chromosome 15. Wu et al. [48] studied the carcass trait of Simmental cattle, and identified the genes in the beef cattle genome significantly associated with foreshank weight and triglyceride levels. A total of 12 and 7 SNPs in the bovine genome were significantly associated with foreshank weight and triglyceride levels, respectively.

In concordance analysis of exterior traits, majorly the QTLs were associated with udder traits (udder swelling score QTL, udder depth QTL, udder attachment QTL, teat length QTL etc.). This information of genotypes could be used to associate phenotypes and perform the selection. Based on the above results, we can assumed that exterior traits are less important for association of QTL with LD block or haplotypes due to insufficient size of QTL and low proportion of concordant QTL with LD blocks. van den Berg et al. [49] studied the concordance for a leg conformation trait in dairy cattle and QTL status was used in a concordance analysis to reduce the number of candidate mutations. In the concordance study of health trait, QTLs associated with somatic cell count were observed almost on every chromosome. The larger size QTL of cold tolerance was observed on chromosome seven. Higher numbers of QTLs associated with Bovine tuberculosis susceptibility were found on chromosome 20 and QTLs for clinical mastitis found on chromosome 14 as well as on chromosome 24. Raphaka et al. [50] identified the markers associated with tuberculosis on *Bos taurus* autosomes (BTA) 2 and on BTA 23 and concluded a major role of BTA 23 for susceptibility to bovine Tuberculosis.

## Conclusion

The study of population structure analysis in Indian buffalo based on SNPs revealed that the distribution of SNP markers across the buffalo genome of all breeds studied was almost similar. Minor differences were observed in various genetic parameters (H_E_, H_O_, F_IS_, F_ST_). The levels of SNPs variation in this study could be insufficient to differentiate the other local breed except Pandharpuri and Jaffarabadi (phenotypically distinct breeds), so there is a need to develop SNP chip based on SNP markers identified by sequence information of local breeds. LD block-QTLs concordance study could explore a new window for genomic selection in animals.

The cattle genome-based SNP information (UMD_3.1) does not offer an optimal coverage for buffalo genome, thereafter the development of new SNP chip based on information of buffalo genome and buffalo-specific genetic technologies is warranted.

## Supporting information captions

**S1 Table: Chromosome-wise distribution of LD-blocks, markers and QTLs for respective Traits**

## References

1. Baker C, Manwell C (1980) Chemical classification of cattle. 1. Breed groups. Animal Blood Groups and Biochemical Genetics 11: 127–150.

2. Barendse W, Harrison BE, Bunch RJ, Thomas MB, Turner LB (2009) Genome wide signatures of positive selection: the comparison of independent samples and the identification of regions associated to traits. BMC genomics 10: 1.

3. Gouveia JJdS, Silva MVGBd, Paiva SR, Oliveira SMPd (2014) Identification of selection signatures in livestock species. Genetics and molecular biology 37: 330–342.

4. Department of Animal Husbandry, Dairying and Fisheries, Govt. of India (2015) Annual Report 2015-16.

5. Food and Agriculural Organisation, UN (2007) The state of the world’s Animal Genetic Reources for Food and Ariculture.

6. Purcell S, Neale B, Todd-Brown K, Thomas L, Ferreira MA, Bender D, et al. (2007) PLINK: a tool set for whole-genome association and population-based linkage analyses. Am J Hum Genet 81: 559–575.

7. Purcell S, Neale B, Todd-Brown K, Thomas L, Ferreira MA, Bender D, et al. (2007) PLINK: a tool set for whole-genome association and population-based linkage analyses. The American Journal of Human Genetics 81: 559–575.

8. Wickham H (2009) ggplot2: Elegant Graphics for Data Analysis.

9. Plotree D, Plotgram D (1989) PHYLIP-phylogeny inference package (version 3.2). Cladistics 5: 6.

10. Barbato M, Orozco-terWengel P, Tapio M, Bruford MW (2015) SNeP: a tool to estimate trends in recent effective population size trajectories using genome-wide SNP data. Frontiers in Genetics 6.

11. Purcell S, Neale B, Todd-Brown K, Thomas L, Ferreira MA, Bender D, et al. (2007) PLINK: a tool set for whole-genome association and population-based linkage analyses. American Journal of Human Genetics 81: 559–575.

12. Ligges U, Mächler M (2003) Scatterplot3d - an R Package for Visualizing Multivariate Data. Journal of Statistical Software. Journal of Statistical Software 8: 1–20.

13. Raj A, Stephens M, Pritchard JK (2014) fastSTRUCTURE: variational inference of population structure in large SNP data sets. Genetics 197: 573–589.

14. Barrett JC, Fry B, Maller J, Daly MJ (2004) Haploview: analysis and visualization of LD and haplotype maps. Bioinformatics 21: 263–265.

15. Patil N, Berno AJ, Hinds DA, Barrett WA, Doshi JM, Hacker CR, et al. (2001) Blocks of limited haplotype diversity revealed by high-resolution scanning of human chromosome 21. Science 294: 1719–1723.

16. Zhang K, Sun F, Waterman MS, Chen T. Dynamic programming algorithms for haplotype block partitioning: applications to human chromosome 21 haplotype data; 2003. ACM. pp. 332–340.

17. Zhang K, Deng M, Chen T, Waterman MS, Sun F (2002) A dynamic programming algorithm for haplotype block partitioning. Proceedings of the National Academy of Sciences 99: 7335–7339.

18. Anderson EC, Novembre J (2003) Finding haplotype block boundaries by using the minimum-description-length principle. The American Journal of Human Genetics 73: 336–354.

19. Gabriel S, Schaffner S, Nguyen H, Moore J, Roy J, Blumenstiel B (2002) The structure of haplotype blocks in the human genome. Science 296.

20. Qin ZS, Niu T, Liu JS (2002) Partition-ligation–expectation-maximization algorithm for haplotype inference with single-nucleotide polymorphisms. The American Journal of Human Genetics 71: 1242–1247.

21. Hu ZL, Park CA, Wu XL, Reecy JM (2013) Animal QTLdb: an improved database tool for livestock animal QTL/association data dissemination in the post-genome era. Nucleic Acids Research 41: D871–879.

22. Quinlan AR, Hall IM (2010) BEDTools: a flexible suite of utilities for comparing genomic features. Bioinformatics 26: 841–842.

23. Kataria R, Sunder S, Malik G, Mukesh M, Kathiravan P, Mishra B (2009) Genetic diversity and bottleneck analysis of Nagpuri buffalo breed of India based on microsatellite data. Russian journal of genetics 45: 826–832.

24. Joshi J, Salar R, Banerjee P (2013) Genetic Variation and Phylogenetic Relationships of Indian Buffaloes of Uttar Pradesh. Asian-Australasian Journal of Animal Sciences 26: 1229.

25. Kumar S, Gupta J, Kumar N, Dikshit K, Navani N, Jain P, et al. (2006) Genetic variation and relationships among eight Indian riverine buffalo breeds. Molecular Ecology 15: 593–600.

26. Joshi J, Salar R, Banerjee P, Sharma U, Tantia M, Vijh R (2015) Assessment of Genetic Variability and Structuring of Riverine Buffalo Population (Bubalus bubalis) of Indo-Gangetic Basin. Animal Biotechnology 26: 148–155.

27. Tantia M, Vijh R, Mishra B, Kumar S, Arora R (2006) Multilocus genotyping to study population structure in three buffalo populations of India. ASIAN AUSTRALASIAN JOURNAL OF ANIMAL SCIENCES 19: 1071.

28. Pundir R, Sahana G, Navani N, Jain P, Singh D, Kumar S, et al. (2000) Characterization of Mehsana buffaloes in India. Animal Genetic Resources 28: 53–62.

29. Dash S, Singh A, Bhatia A, Jayakumar S, Sharma A, Singh S, et al. (2017) Evaluation of Bovine High-Density SNP Genotyping Array in Indigenous Dairy Cattle Breeds. Animal Biotechnology: 1–7.

30. Iamartino D, Williams JL, Sonstegard T, Reecy J, Tassell Cv, Nicolazzi EL, et al. (2013) The buffalo genome and the application of genomics in animal management and improvement. Buffalo Bulletin 32: 151–158.

31. MacEachern S, Hayes B, McEwan J, Goddard M (2009) An examination of positive selection and changing effective population size in Angus and Holstein cattle populations (Bos taurus) using a high density SNP genotyping platform and the contribution of ancient polymorphism to genomic diversity in Domestic cattle. BMC Genomics 10: 181.

32. Dadi H, Kim JJ, Kim KS, Yoon D (2012) Evaluation of Single Nucleotide Polymorphisms (SNPs) Genotyped by the Illumina Bovine SNP50K in Cattle Focusing on Hanwoo Breed. Asian-Australasian Journal of Animal Sciences 25: 28–32.

33. Lango-Allen H, Estrada K, Lettre G, Berndt SI, Weedon MN, Rivadeneira F, et al. (2010) Hundreds of variants clustered in genomic loci and biological pathways affect human height. Nature 467: 832–838.

34. Edea Z, Bhuiyan MS, Dessie T, Rothschild MF, Dadi H, Kim KS (2015) Genome-wide genetic diversity, population structure and admixture analysis in African and Asian cattle breeds. Animal 9: 218–226.

35. Kim ES, Sonstegard TS, Rothschild MF (2015) Recent artificial selection in U.S. Jersey cattle impacts autozygosity levels of specific genomic regions. BMC Genomics 16: 302.

36. McKay SD, Schnabel RD, Murdoch BM, Matukumalli LK, Aerts J, Coppieters W, et al. (2008) An assessment of population structure in eight breeds of cattle using a whole genome SNP panel. BMC Genetics 9: 37.

37. Sraphet S, Moolmuang B, Na-Chiangmai A, Panyim S, Smith DR, Triwitayakorn K (2008) Use of cattle microsatellite markers to assess genetic diversity of Thai Swamp buffalo (Bubalus bubalis). Asian-Australasian Journal of Animal Sciences 21: 177.

38. Suh S, Kim Y-S, Cho C-Y, Byun M-J, Choi S-B, Ko Y-G, et al. (2014) Assessment of genetic diversity, relationships and structure among Korean native cattle breeds using microsatellite markers. Asian-Australasian Journal of Animal Sciences 27: 1548.

39. Machado MA, Schuster I, Martinez ML, Campos AL (2003) Genetic diversity of four cattle breeds using microsatellite markers. Revista Brasileira de zootecnia 32: 93–98.

40. Melka MG, Schenkel FS (2012) Analysis of genetic diversity in Brown Swiss, Jersey and Holstein populations using genome-wide single nucleotide polymorphism markers. BMC Research Notes 5: 161.

41. Shah TM, Patel JS, Bhong CD, Doiphode A, Umrikar UD, Parmar SS, et al. (2013) Evaluation of genetic diversity and population structure of west-central Indian cattle breeds. Animal Genetics 44: 442–445.

42. Kumar S, Nagarajan M, Sandhu JS, Kumar N, Behl V (2007) Phylogeography and domestication of Indian river buffalo. BMC Evolutionary Biology 7: 186.

43. Shin DH, Cho KH, Park KD, Lee HJ, Kim H (2013) Accurate Estimation of Effective Population Size in the Korean Dairy Cattle Based on Linkage Disequilibrium Corrected by Genomic Relationship Matrix. Asian-Australasian Journal of Animal Sciences 26: 1672–1679.

44. Santana M, Aspilcueta-Borquis R, Bignardi A, Albuquerque LG, Tonhati H (2011) Population structure and effects of inbreeding on milk yield and quality of Murrah buffaloes. Journal of Dairy Science 94: 5204–5211.

45. Villa-Angulo R, Matukumalli LK, Gill CA, Choi J, Van Tassell CP, Grefenstette JJ (2009) High-resolution haplotype block structure in the cattle genome. BMC Genetics 10: 19.

46. Jiang L, Liu J, Sun D, Ma P, Ding X, Yu Y, et al. (2010) Genome wide association studies for milk production traits in Chinese Holstein population. PloS One 5: e13661.

47. Mai M, Sahana G, Christiansen F, Guldbrandtsen B (2010) A genome-wide association study for milk production traits in Danish Jersey cattle using a 50K single nucleotide polymorphism chip. Journal of animal science 88: 3522–3528.

48. Wu Y, Fan H, Wang Y, Zhang L, Gao X, Chen Y, et al. (2014) Genome-wide association studies using haplotypes and individual SNPs in Simmental cattle. PLoS One 9: e109330.

49. van den Berg I, Fritz S, Rodriguez S, Rocha D, Boussaha M, Lund MS, et al. (2014) Concordance analysis for QTL detection in dairy cattle: a case study of leg morphology. Genetics, Selection, Evolution 46: 31.

50. Raphaka K, Matika O, Sánchez-Molano E, Mrode R, Coffey MP, Riggio V, et al. (2017) Genomic regions underlying susceptibility to bovine tuberculosis in Holstein-Friesian cattle. BMC Genetics 18: 27.

